# Bispecific Targeting of CHI3L1 and PD-1/PD-L1 Axis as a Novel Therapeutic Strategy for Idiopathic Pulmonary Fibrosis

**DOI:** 10.1101/2025.09.17.676854

**Authors:** Han-Seok Jeong, Takayuki Sadanaga, Joyce H Lee, Suchitra Kamle, Bing Ma, Yang Zhou, Sung Jae Shin, Jack A. Elias, Chun Geun Lee

## Abstract

CHI3L1, a chitinase-like protein, plays a key role in the pathogenesis of pulmonary fibrosis, though the precise mechanisms remain unclear. This study explores how CHI3L1 regulates profibrotic macrophage activation and invasive myofibroblast differentiation and their interactions. *In vitro*, CHI3L1 induced profibrotic M2 macrophage activation and differentiation marked by increased expression of CD163, CD206, and PD-L1. CHI3L1 also enhanced TGF-β_1_ effects on lung fibroblasts including myofibroblast transformation, migration and tissue invasion. Mechanistically, CHI3L1 increased TGF-β_1_-stimulation of Smad, Akt and Erk signaling and PD-L1 played a significant role in TGF-β_1_/CHI3L1-stimulated myofibroblast transformation. Coculture experiment further confirmed the ability of CHI3L1 to induce profibrotic macrophage activation that enhanced myofibroblast transformation mediated via a CD44-PD-L1 axis. Following *in vivo* bleomycin challenge, CHI3L1 transgenic mice exhibited significantly higher levels of PD-L1^+^ M2 macrophages, PD-L1^+^/PDGFRα^+^ fibroblasts and increased numbers of PD-1^+^ and CD45^+^/PD-1^+^ cells compared to wild-type controls. Notably, combined treatment with anti-CHI3L1 and anti-PD-1 antibodies, or a bispecific anti-CHI3L1-anti-PD-1 antibody, resulted in greater inhibition of bleomycin-induced fibrosis than either antibody alone. These findings suggest that there is a stimulatory interaction between CHI3L1 and the PD-1/PD-L1 axis in promoting profibrotic macrophage activation and invasive fibroblast differentiation. The results also highlight the potential of bispecific targeting of CHI3L1 and the PD-1/PD-L1 pathway as an effective therapeutic approach for pulmonary fibrosis.

## Introduction

Pulmonary fibrosis (PF) is a progressive and often fatal lung disease characterized by excessive deposition of extracellular matrix (ECM) proteins, leading to scarring and compromised lung function [1]. Idiopathic pulmonary fibrosis (IPF) is the most common form of pulmonary fibrosis and has a median survival of only 2-3 years after diagnosis [2, 3]. The pathogenesis of PF involves complex interactions between various cell types, including fibroblasts, myofibroblasts, and immune cells such as macrophages [4, 5]. These cellular interactions drive a self-sustaining fibrotic cycle, marked by the activation and differentiation of macrophages and fibroblasts which contribute to the perpetuation of fibrotic tissue remodeling [6, 7]. Recent research has highlighted chitinase-3-like protein 1 (CHI3L1) as a crucial regulator in this setting [8, 9]. However, the precise mechanisms through which CHI3L1 drives fibrosis remain incompletely understood.

The glycosyl hydrolase gene family 18 (GH18) contains chitinases and chitinase-like proteins (CLPs) that lack enzyme activity. CHI3L1 (also known as YKL-40 in humans and *Chil1* in mice), the prototypic CLP, has been implicated in multiple biologic processes including cell proliferation, migration, inflammation, and tissue remodeling associated with various diseases in the lung and other organs [10–14]. Elevated levels of CHI3L1 have been observed in patients with idiopathic pulmonary fibrosis (IPF) which correlate with disease severity. These studies suggest that CHI3L1 contributes to lung fibrosis by modulating immune responses and/or fibroblast activation [8, 9, 15]. However, the mechanisms by which CHI3L1 promotes fibrotic pathways remain incompletely understood, particularly in the context of macrophage activation and myofibroblast differentiation.

Macrophages have recently been recognized as central players in the fibrotic response, with plasticity to adopt pro-inflammatory (M1) or anti-inflammatory and profibrotic (M2) phenotypes in response to environmental stimuli [16, 17]. The M2 phenotype is characterized by the expression of markers such as CD163 and CD206 and supports tissue repair and fibrosis through the secretion of growth factors and cytokines that promote fibroblast proliferation and activation [18, 19]. Studies from our laboratory and others have shown that CHI3L1 promotes the differentiation of monocytes into M2-like macrophages [15, 20], thereby contributing to the profibrotic milieu. Interestingly, we recently observed that CHI3L1 is also a powerful stimulator of PD-L1 [21], an immune checkpoint molecule associated with immune evasion and chronic inflammation [22, 23]. Surprisingly, the mechanisms underlying these responses have not been fully defined.

The role of the PD-1/PD-L1 axis in fibrotic diseases has gained interest, as it facilitates immune tolerance and promotes profibrotic responses [24, 25]. In the context of pulmonary fibrosis, PD-L1 is implicated in the development of invasive fibroblasts responsible for persistent and progressive pulmonary fibrosis [26]. Although CHI3L1 is a strong stimulator of PD-1/PD-L1 expression in immune and structural cells including macrophages, fibroblasts and epithelial cells in the tumor microenvironment [21], the mechanistic details of CHI3L1-driven macrophage differentiation and fibroblast activation and their coordination with the regulated expression of PD-1/PD-L1 require further investigation.

In this study, we investigated the roles of CHI3L1 in regulating profibrotic macrophage activation and invasive myofibroblast differentiation in the pathogenesis of pulmonary fibrosis. Through *in vitro* and *in vivo* experiments, we explored the interactions between CHI3L1, CD44, PD-L1, and key profibrotic pathways, including TGF-β signaling. Additionally, we assessed the therapeutic potential of bispecific targeting of CHI3L1 and PD-1/PD-L1 axis in a murine model of bleomycin-induced pulmonary fibrosis. Our findings provide novel mechanistic insights into the pathogenesis of pulmonary fibrosis and underscore the therapeutic promise of dual targeting of CHI3L1 and the PD-1/PD-L1 axis.

## Results

### IL-13 and TGF-β_1_ stimulate profibrotic macrophage activation via a CHI3L1-dependent mechanism

To investigate the role of CHI3L1 in macrophage activation, we prepared bone marrow-derived macrophages (BMDMs) from WT and CHI3L1-null mutant (*Chil1*^-/-^) mice and stimulated them with recombinant IL-13, TGF-β_1_, and IFN-γ. FACS analysis showed that recombinant IL-13 and TGF-β stimulation alone or together increased the frequency of CD163^+^ or CD206^+^ macrophages in WT mice, whereas these increases were significantly reduced in CHI3L1-null mice (Fig. 1A and Supplemental Fig. S1). Conversely, IFN-γ stimulation did not substantially increase the frequency of CD163^+^ and CD206^+^ macrophages, and its effects were not significantly altered by null mutations of CHI3L1 (Fig. 1A and Supplemental Fig. S1). Similarly, CX3CR1^+^/SiglecF^+^ macrophages were increased when treated with IL-13 and TGF-β_1_, but not with IFN-γ, and these increases were absent in CHI3L1-null mice (Fig. 1B and Supplemental Fig. S1). These results demonstrate that IL-13 and TGF-β_1_, alone and in combination, stimulate profibrotic M2 macrophage activation via a CHI3L1-dependent mechanism.

**Figure 1.**
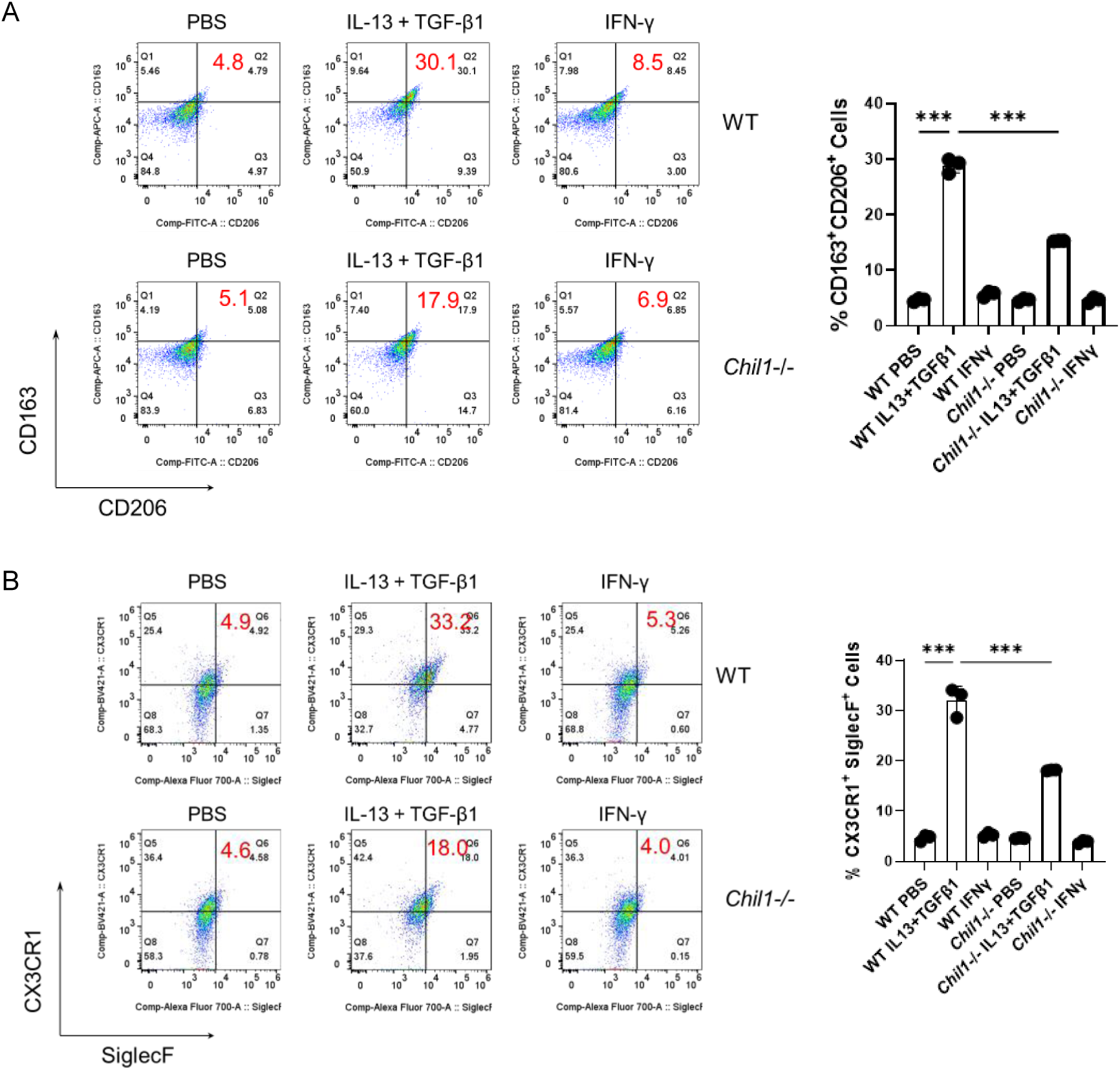
IL-13 and TGF-β_1_ stimulate M2 macrophage polarization via a CHI3L1- dependent mechanism. Bone marrow-derived macrophages (BMDMs) were isolated from wild-type (WT) and CHI3L1-null (*Chil1*^-/-^) mice and stimulated with recombinant IL-13 (20 ng/mL), TGF-β_1_ (5 ng/mL), and IFN-γ (20 ng/mL) for 72 hours. Cells were then analyzed by flow cytometry. (A) Frequency of CD206^+^/CD163^+^ M2 macrophages. (B) Frequency of CX3CR1^+^/SiglecF^+^ profibrotic macrophages. The bar graphs show the mean ± SEM from biological replicates. \*\*\**p* < 0.001 by one-way ANOVA with multiple comparisons.

### Recombinant CHI3L1 induces profibrotic M2 macrophage differentiation and PD-L1 expression

To determine whether CHI3L1 drives profibrotic M2 macrophage differentiation, we evaluated the effect of CHI3L1 on the macrophage differentiation using THP-1 monocytic cells. Recombinant CHI3L1 treatment increased the frequency of CD206^+^/CD163^+^ macrophages and this CHI3L1 effect was blocked by either anti-CHI3L1 monoclonal antibody (referred to as FRG antibody) or kasugamycin (KSM), a pan-chitinase inhibitor (Fig. 2A). Since studies from our laboratory and others identified PD-L1 (*CD274*) as a critical mediator of CHI3L1- induced immune tolerance and alternative macrophage activation [21, 27], we further assessed PD-L1 expression in CHI3L1-stimulated macrophages. PD-L1 expression was significantly increased in the differentiated macrophages with CHI3L1 stimulation (similarly to CD163 and CD206) at both the protein and mRNA levels, and these increases were abrogated by FRG or KSM treatment (Fig. 2, B and C). These findings demonstrate that CHI3L1 can directly stimulate CD206^+^/CD163^+^ profibrotic M2 macrophage differentiation with increased expression of PD-L1.

**Figure 2.**
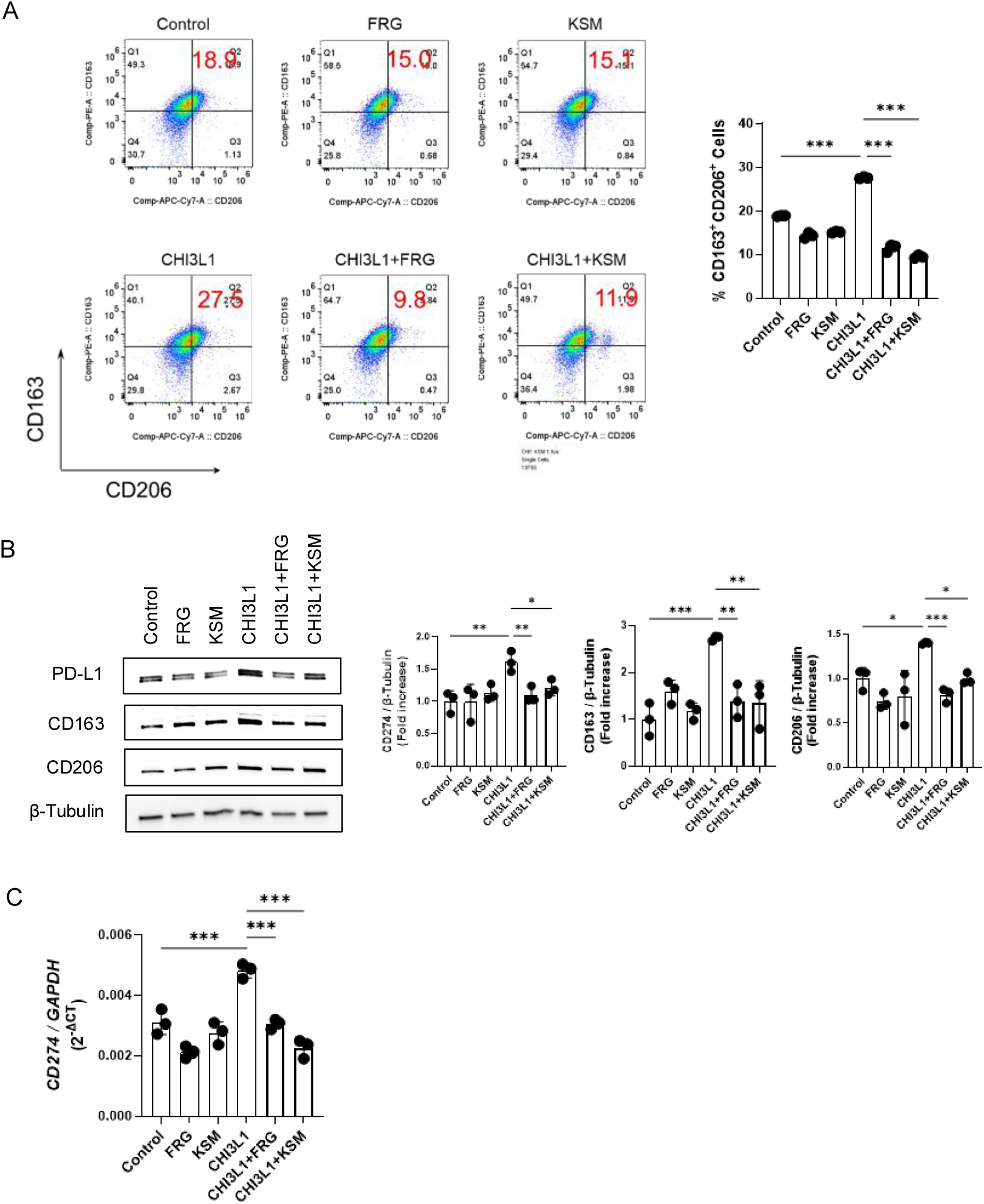
CHI3L1 drives M2 macrophage differentiation and PD-L1 expression. Human monocytic THP-1 cells were stimulated with phorbol 12-myristate 13-acetate (PMA, 100 ng/mL) for 24 hours followed by 24 hours of rest and then stimulated with recombinant CHI3L1 (500 ng/mL) in the presence of anti-CHI3L1 antibody (FRG, 250 ng/mL) or pan-chitinase inhibitor kasugamycin (KSM, 250 ng/mL) for 24 hours. Expression of PD-L1, CD206, and CD163 was then evaluated. (A) Flow cytometric analysis of the CD206^+^/CD163^+^ M2 macrophages. (B) Western blot analysis of CD163, CD206, and PD-L1 protein expression. β-tubulin was used as a loading control. (C) *CD274* (PD-L1) mRNA expression measured by quantitative RT-PCR. The bar graphs in panel B represent densitometric quantification of Western blot signals; values in panels A-C represent the mean ± SEM. \**p* < 0.05, \*\**p* < 0.01, \*\*\**p* < 0.001 by one-way ANOVA with multiple comparisons.

### CHI3L1 enhances TGF-β_1_-stimulated myofibroblast transformation, migration, and invasiveness

The role of CHI3L1 in fibroblastto-myofibroblast transformation was assessed in normal human lung fibroblasts (NHLF) treated with recombinant CHI3L1, TGF-β_1_, or both. Immunocytochemical analyses and Western blot analyses revealed minimal increases in α-SMA expression in response to CHI3L1 stimulation alone, while maximal induction was observed in the presence of both CHI3L1 and TGF-β_1_ (Fig. 3A and B). Similarly, CHI3L1 alone did not affect fibroblast migration and invasion, but it significantly enhanced TGF-β_1_- induced migration and invasion with maximal effects being observed in simultaneous stimulation with CHI3L1 and TGF-β_1_ (Fig. 3, C and D). All these responses were attenuated by either anti-CHI3L1 antibody (FRG) or KSM treatment, indicating a specific contribution of CHI3L1 to TGF-β_1_-driven myofibroblast transformation, migration, and invasiveness of lung fibroblasts.

**Figure 3.**
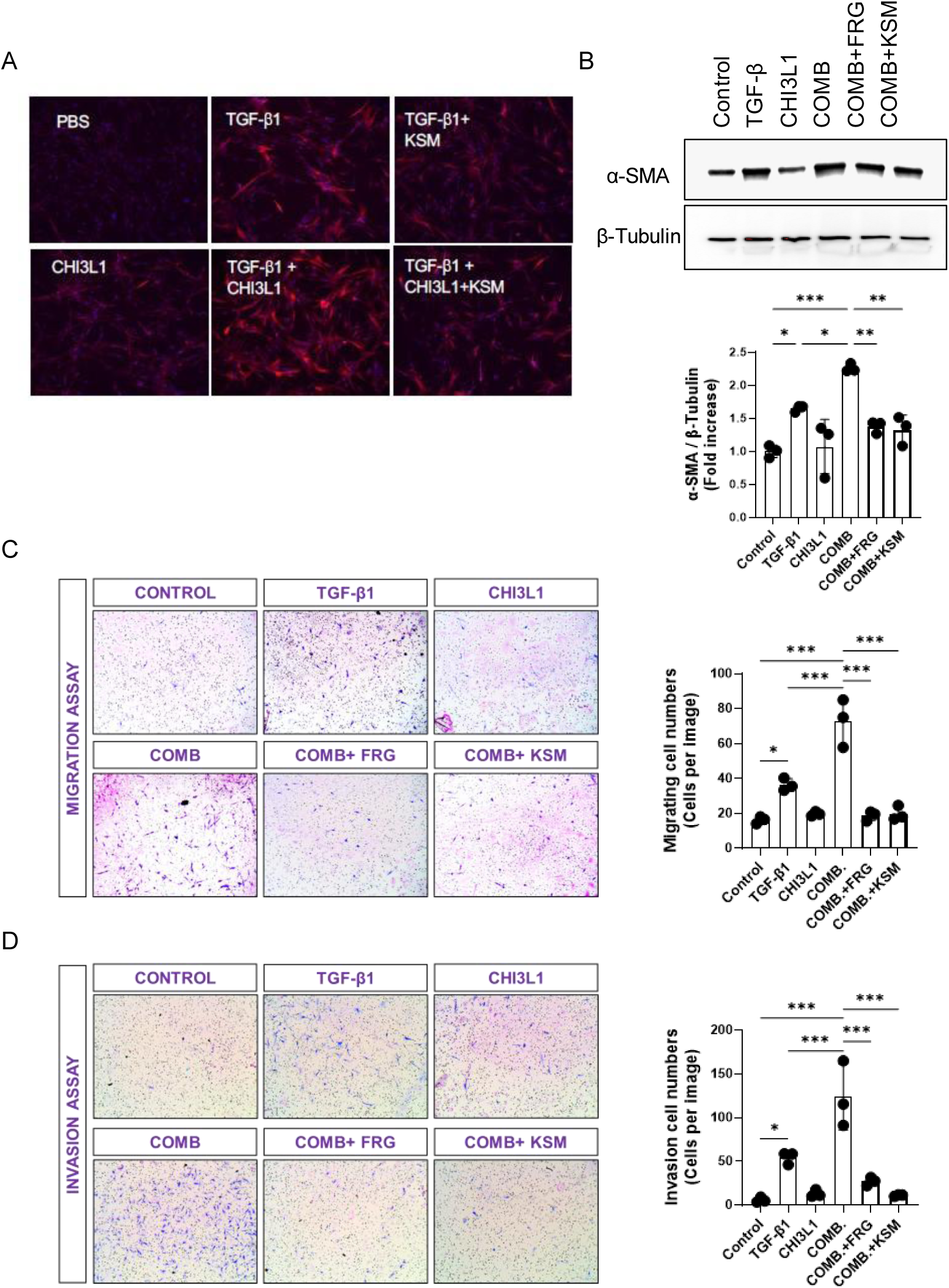
CHI3L1 enhances TGF-β_1_–induced myofibroblast transformation, migration, and invasion. Normal human lung fibroblasts (NHLFs) were treated with recombinant CHI3L1 (500 ng/mL) and/or TGF-β_1_ (5 ng/mL) in the presence or absence of an anti-CHI3L1 antibody (FRG, 250 ng/mL) or the chitinase inhibitor KSM. (A-B) α-SMA expression was assessed by immunohistochemistry (IHC) and Western blot analysis. (C-D) Fibroblast migration and invasion assays were performed using 24 transwell culture (8 μm pore size) plates. In each the number of cells that migrated or invaded through the membrane were counted. The bar graph in panel B represents the densitometric quantification of Western blot signals. β-tubulin was used as a loading control. The values in panels B and C represent the mean ± SEM. \**p* < 0.05, \*\**p* < 0.01, \*\*\**p* < 0.001 by one-way ANOVA, multiple comparisons. COMB, combined treatment with CHI3L1 and TGF-β_1_.

### CHI3L1 enhances TGF-β_1_ signaling and its effector functions in lung fibroblasts through the CD44-PD-L1 axis

To dissect the mechanistic interplay between CHI3L1 and TGF-β_1_ signaling in lung fibroblasts, we first examined the impact of recombinant CHI3L1 on TGF-β_1_-induced signaling in normal human or mouse lung fibroblasts. CHI3L1 augmented TGF-β_1_-induced activation of SMAD2 (p-SMAD2), ERK (p-ERK), and AKT (p-AKT), and these effects were abrogated by treatment with FRG antibody or KSM (Fig. 4A). Given that CD44 acts as a co-receptor for CHI3L1 and its involvement in TGF-β_1_ signaling [28, 29], we investigated whether the CHI3L1-TGF-β_1_ interaction is mediated through CD44 using both control and CD44 knockdown fibroblasts. As shown in Fig. 4B, the effects of CHI3L1 alone and the interaction of CHI3L1 with TGF-β_1_- induced ERK activation was markedly diminished in CD44-silenced mouse lung fibroblasts. In contrast, no significant changes were observed in CHI3L1 induced SMAD2 or AKT activation (Fig. 4B). To further define the downstream functional consequences, we evaluated α-smooth muscle actin (α-SMA) expression as a marker of myofibroblast transformation. CD44 knockdown abrogated the synergistic effect of CHI3L1 and TGF-β on α-SMA and PD-L1 induction (Fig. 4C). Moreover, PD-L1 knockdown similarly attenuated CHI3L1/TGF-β1- induced α-SMA expression, while siRNA control had no effect (Fig. 4D). CD44 neutralization using an anti-CD44 antibody similarly inhibited the stimulatory interaction between CHI3L1 and TGF-β_1_ on PD-L1 and α-SMA expression (Supplemental Fig. S2). Collectively, these results suggest that a CHI3L1-CD44-ERK-PD-L1 signaling axis amplifies TGF-β_1_ responses and promotes profibrotic transformation in lung fibroblasts.

**Figure 4.**
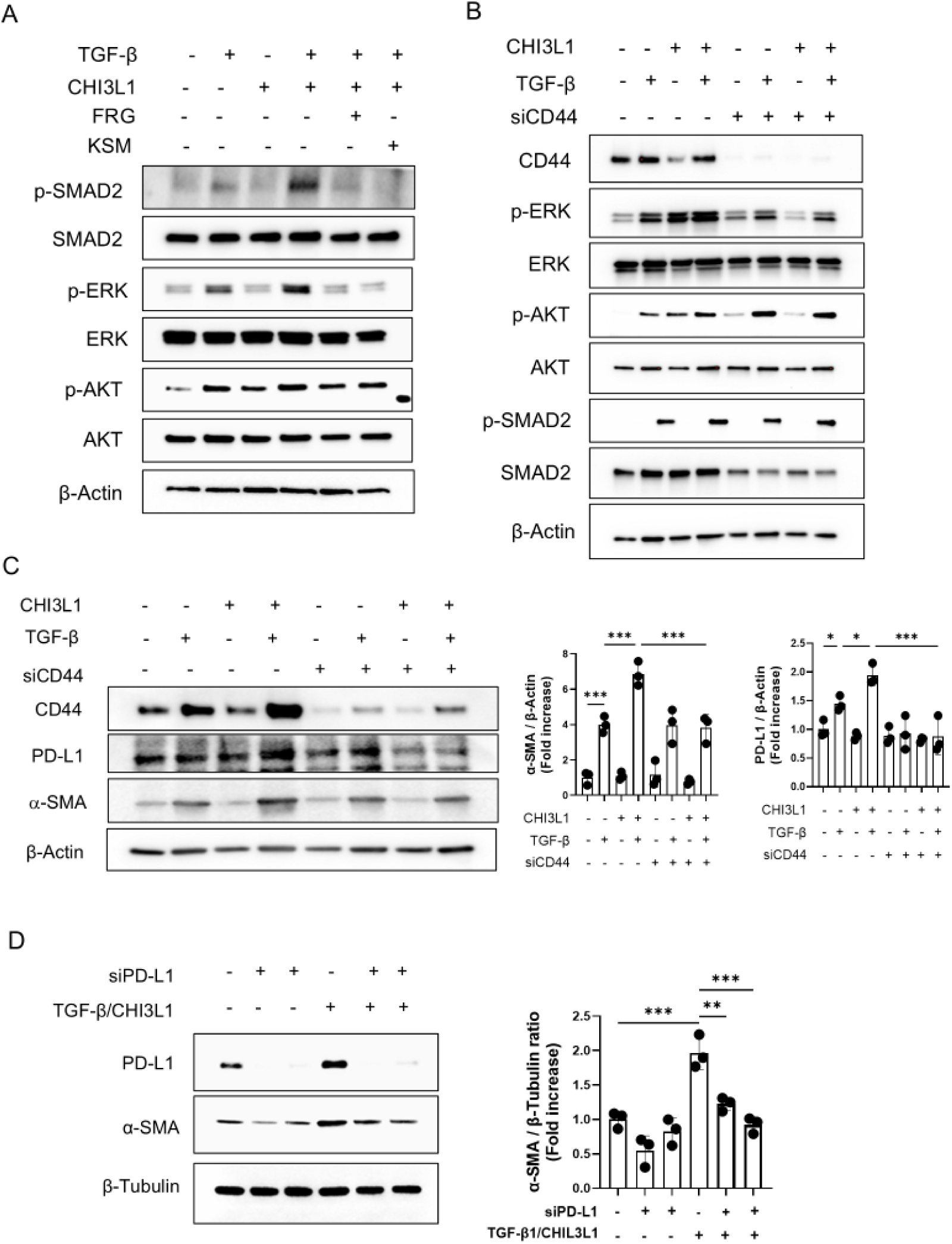
CHI3L1 promotes TGF-β signaling and myofibroblast transformation through CD44/PD-L1 axis. NHLFs or mouse lung fibroblasts were treated with recombinant CHI3L1 (500 ng/mL) and/or TGF-β_1_ (5 ng/mL) in the presence or absence of chitinase inhibitor an anti-CHI3L1 antibody (FRG, 250 ng/mL), or KSM (250 ng/mL). (A) Western blot analysis of SMAD, AKT, and ERK phosphorylation in human lung fibroblasts treated with CHI3L1 and/or TGF-β_1_. (B) Western blot analysis of the same signaling pathways in mouse lung fibroblasts treated with or without CD44 siRNA (siCD44). (C) Western blot analysis of PD-L1 and α-SMA expression in lung fibroblasts stimulated with CHI3L1 and/or TGF-β in the presence or absence of CD44 silencing. (D) Western blot analysis of α-SMA expression in NHLFs with or without PD-L1 silencing to assess the role of PD-L1 in myofibroblast transformation. β-actin was used as a loading control. Panels A and B show representative results from three independent experiments. Bar graphs in panels C and D represent densitometric quantification of protein expression (mean ± SEM). \**p* < 0.05, \*\**p* < 0.01, \*\*\**p* < 0.001 by one-way ANOVA with multiple comparisons. COMB, combined treatment with CHI3L1 and TGF-β.

### CHI3L1 facilitates profibrotic macrophage-fibroblast crosstalk via the CD44-PD-L1 axis

Profibrotic macrophage polarization and aberrant fibroblast activation are key contributors to pulmonary fibrosis [30]. To investigate the role of CHI3L1 in macrophage–fibroblast communication, we performed co-culture experiments using macrophages from wild-type (WT) and CHI3L1-deficient (*Chil1*^-/-^) mice and fibroblasts with or without CD44 silencing, as outlined schematically in Fig. 5A and 5C. In the first set of experiments, normal fibroblasts were co-cultured with WT or CHI3L1-null macrophages that had been pre-stimulated with IL-13 and TGF-β_1_ to induce a profibrotic phenotype (Fig. 5A). The fibroblast response was assessed by measuring α-smooth muscle actin (α-SMA) and PD-L1 expression. IL-13/TGF-β_1_- stimulated WT macrophages significantly induced both α-SMA and PD-L1 expression in fibroblasts, whereas CHI3L1-null macrophages failed to do so (Fig. 5B). In additional experiments, fibroblasts isolated from WT mice, with or without CD44 knockdown, were co-cultured with IL-13/TGF-β_1_-stimulated WT macrophages (Fig. 5C). CD44-silenced fibroblasts exhibited reduced expression of PD-L1 and α-SMA compared to control fibroblasts (Fig 5C). Taken together, these results demonstrate that the interactive stimulation caused by CHI3L1 and TGF-β_1_ on PD-L1^+^ myofibroblast activation is mediated through CD44, highlighting CD44-PD-L1 axis as a critical pathway in the macrophage-myofibroblast crosstalk.

**Figure 5.**
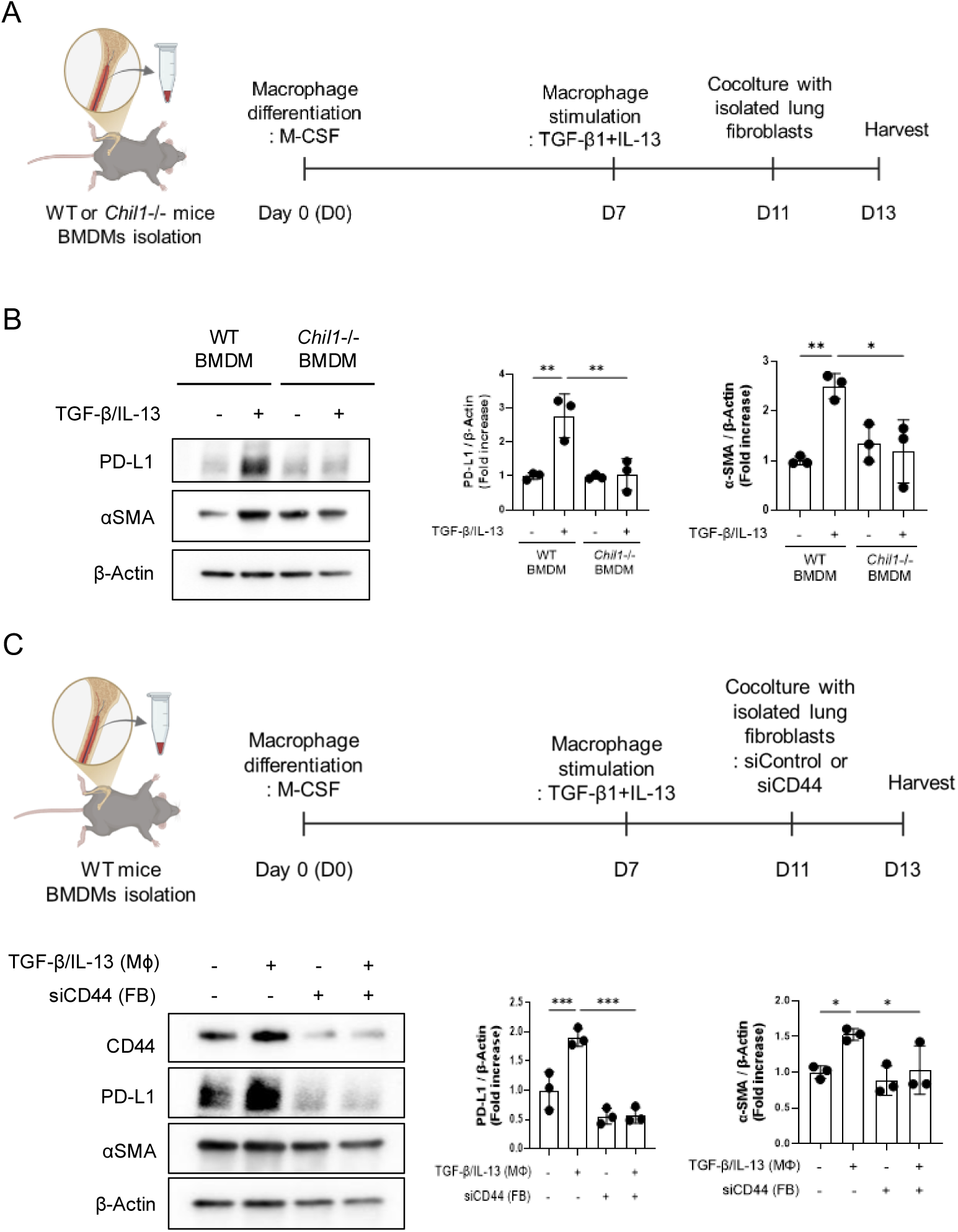
CHI3L1 facilitates profibrotic macrophages-fibroblast crosstalk via the CD44- PD-L1 axis. (A-B) Schematic of the co-culture experiment in which mouse lung fibroblasts were cultured with wild-type (WT) or CHI3L1-deficient (*Chil1*⁻/⁻) macrophages pre-stimulated with IL-13 (20 ng/mL) and TGF-β1 (5 ng/mL) and Western blot analysis of PD-L1 and α-SMA expression in mouse lung fibroblasts. (C-D) Schematic of the co-culture experiment using WT macrophages and fibroblasts with or without CD44 silencing and Western blot analysis of PD-L1 and α-SMA expression in mouse lung fibroblasts. The bar graphs in panels B and D represent densitometric quantification of protein expression (mean ± SEM). \**p* < 0.05, \*\**p* < 0.01, \*\*\**p* < 0.001 by one-way ANOVA, multiple comparisons.

### CHI3L1 increases profibrotic M2 macrophages and invasive fibroblasts in bleomycin-challenged lungs

To evaluate the *in vivo* role of CHI3L1 in driving profibrotic macrophage responses, wild-type (WT) and CHI3L1 overexpressing transgenic (Tg) mice were treated with bleomycin. In WT lungs, bleomycin exposure led to increased accumulation of CD206^+^/CD163^+^ and SiglecF^+^/CX3CR1^+^ profibrotic macrophages, and these responses were further augmented in CHI3L1 Tg mice (Fig. 6A and Supplemental Fig. S3). Notably, the number of PD-L1^+^/CD206^+^ and PD-L1⁺/CD163⁺ macrophages was significantly elevated in CHI3L1 Tg lungs compared to WT (Fig. 6B). Immunohistochemistry (IHC) analysis further confirmed the expansion of CD206^+^/CD163^+^ macrophages and revealed a notable increase in PD-L1^+^/PDGFRα^+^ mesenchymal cells, including fibroblasts, in the lungs of CHI3L1 Tg mice relative to controls (Fig. 6C). These findings indicate that CHI3L1 enhances the development of PD-L1- expressing profibrotic macrophages and invasive fibroblasts in the setting of bleomycin-induced lung injury. In addition, we observed a marked increase in total and CD45⁺/PD-1⁺ immune cells including CD45^+^/CD11c^+^/PD-1^+^ macrophages in the lungs of CHI3L1 Tg mice compared to WT controls (Supplemental Fig. S4), suggesting that enhanced PD-1/PD-L1 interactions contribute to the immunosuppressive and fibrotic microenvironment orchestrated by CHI3L1.

**Figure 6.**
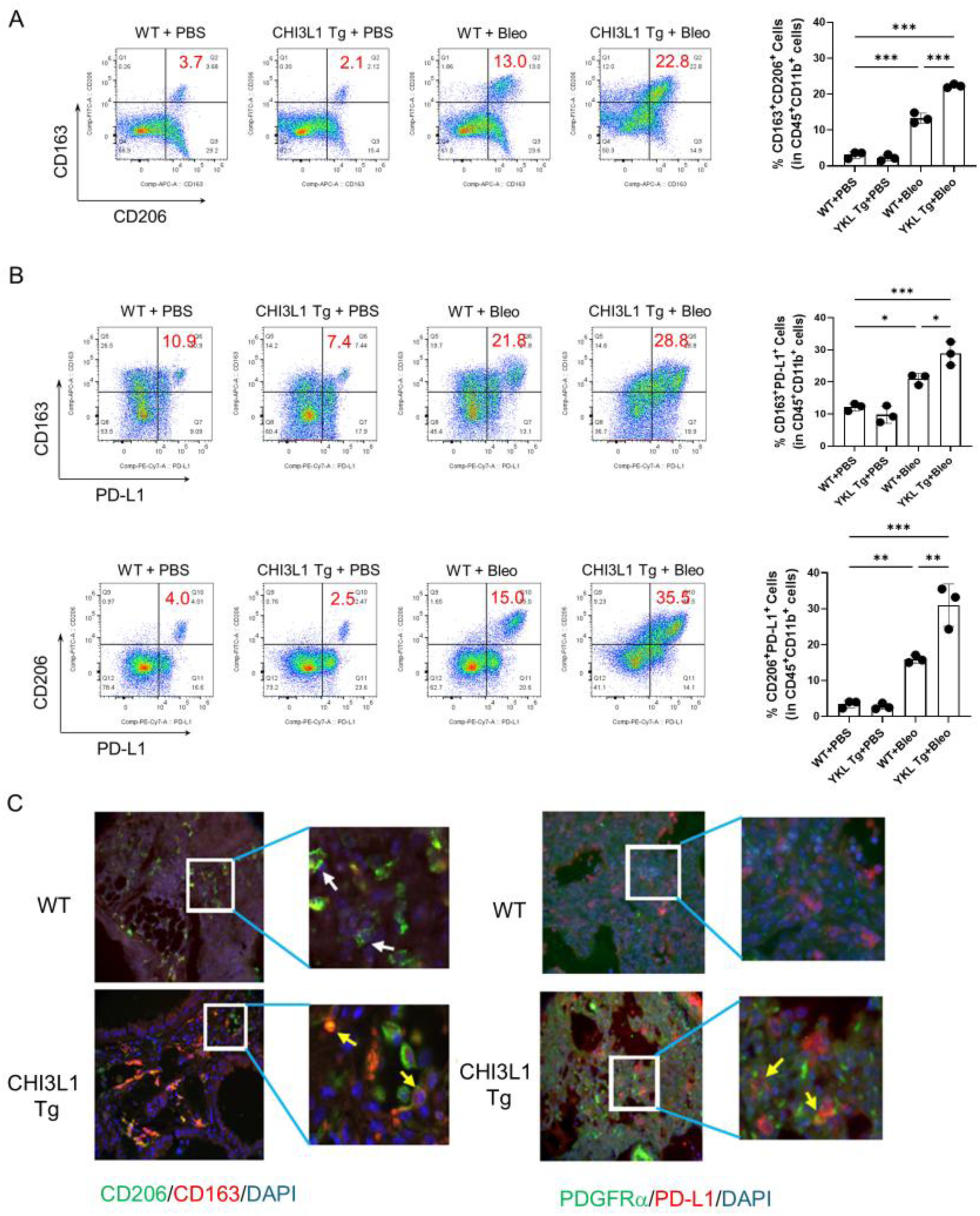
**CHI3L1 increases profibrotic macrophages and fibroblasts in bleomycin-challenged lungs. 6-8 weeks old WT and** CHI3L1 (YKL) Tg mice were challenged with PBS or bleomycin. Mice were sacrificed 14 days after bleomycin challenge and analyzed. (A) Flow cytometric analysis of CD206⁺/CD163⁺ and PD-L1⁺ macrophages in lungs of WT and CHI3L1 Tg mice. (B) Immunohistochemical analysis of CD206⁺/CD163⁺ macrophages and PD-L1^+^/PDGFRα^+^ fibroblasts in WT and CHI3L1 Tg lungs. Flow cytometry and IHC data in panels A-C are representative of at least 3 independent experiments.

### Therapeutic potential of anti-CHI3L1, anti-PD-1, and bispecific (anti-CHI3L1×PD-1) antibodies in bleomycin-induced lung fibrosis

Because our studies demonstrated that CHI3L1 plays a significant role in profibrotic macrophage activation and fibroblast responses through the CD44-PD-1/PD-L1 axis, we next evaluated the *in vivo* therapeutic potential of mono- or bi-specific antibodies targeting CHI3L1 and the PD-1/PD-L1 axis in bleomycin-induced pulmonary fibrosis. Mice treated with bleomycin and antibody control (IgG) showed significant increases in pulmonary collagen content and the expression of *Col1a1*, *Acta2* (the gene for α-SMA), *Cd206,* and *Cx3cr1* (Fig. 7). Importantly, treatment with either the bispecific antibody or a combination of anti-PD-1 and anti-CHI3L1 (FRG) antibodies resulted in the greatest reduction in lung fibrosis, as shown by histological analysis and collagen deposition compared to WT controls (Fig. 7, A and B). Gene expression analysis demonstrated similar reductions in the fibrosis markers (*Col1a1* and *Acta2*) and profibrotic macrophage markers (*Cd206* and *Cx3cr1*) with these treatments (Fig. 7C). FACS analysis corroborated these findings, showing reduced frequency of profibrotic macrophages in mice treated with the bispecific or combination therapies (Supplemental Fig. S5). These studies suggest that the simultaneous targeting of CHI3L1 and the PD-1-PD-L1 axis provides effective anti-fibrotic effects on bleomycin-stimulated lung fibrosis.

**Figure 7.**
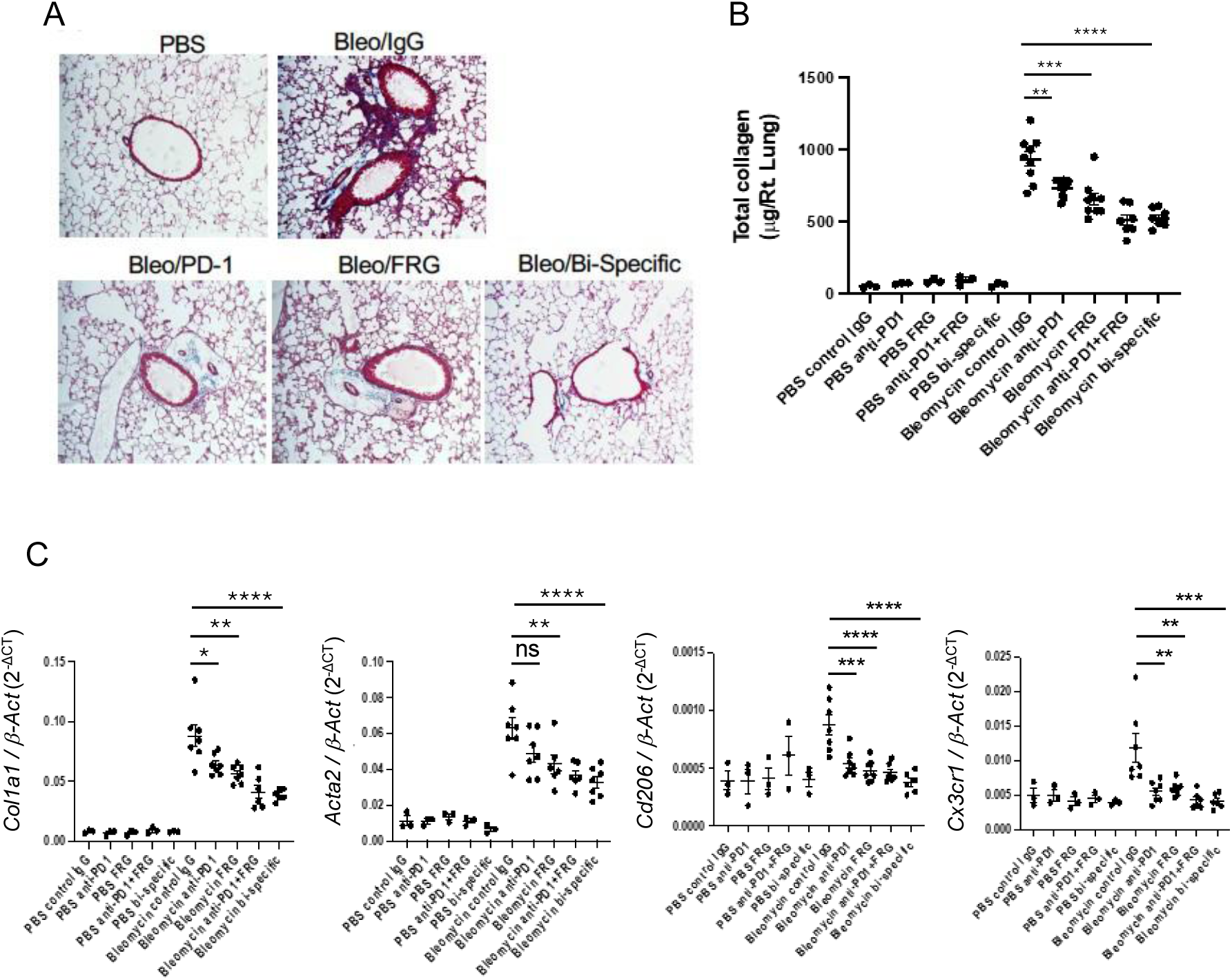
Anti-CHI3L1, anti–PD-1, and bispecific antibodies attenuate bleomycin-induced lung fibrosis. Wild-type (WT) mice (6–8 weeks old) were challenged intratracheally with PBS or bleomycin and subsequently treated with anti-CHI3L1 antibody (FRG), anti–PD-1 antibody, or a bispecific anti-CHI3L1×PD-1 antibody, either individually or in combination. Mice were sacrificed on day 14 post-bleomycin challenge and analyzed for fibrosis. (A) Histological assessment and Masson’s trichrome staining of lung tissues from bleomycin-challenged mice treated with FRG, anti–PD-1, or the bispecific antibody (B) Quantification of total lung collagen content. (C) qRT-PCR analysis of the expression of fibrotic markers (*Col1a1*, *Acta2*) and profibrotic macrophage markers (*Cd206*, *Cx3cr1*). The values in panels B and C are mean ± SEM. \**p* < 0.05, \*\**p* < 0.01, \*\*\**p* < 0.001, \*\*\*\**p* < 0.0001 by one-Way ANOVA with multiple comparisons.

### Increased CHI3L1 and PD-1 expression and spatial interaction between CHI3L1⁺ macrophages and CD44⁺ fibroblasts in lungs of IPF patients compared to controls

To assess the human relevance of our findings, we analyzed publicly available RNA profiling datasets from patients with interstitial lung disease (ILD) and controls. First, we examined a large microarray dataset (NCBI GEO: GSE47460) [31] from the Lung Tissue Research Consortium (LTRC), which includes samples from 254 ILD patients, 220 COPD patients, and 108 controls. Diagnoses were based on clinical history, CT imaging, and surgical pathology. In this dataset, *CHI3L1* and *PDCD1* (PD-1) transcript levels were significantly elevated in the lungs of IPF patients compared to controls (Fig. 8A). Notably, co-expression of *CHI3L1* and *PDCD1* was inversely correlated with pulmonary function indices, including % predicted forced vital capacity (FVC) and diffusion capacity of the lungs for carbon monoxide (DLCO) (Fig. 8B), suggesting functional relevance of these genes in disease outcomes. Moreover, mean *CHI3L1* and *PDCD1* expression progressively increased from healthy controls to IPF patients, with further stratification by disease severity (Fig. 8C). Pairwise analyses showed significant differences between Control vs Stage I (p = 0.016), Stage I vs Stage II+III (p = 0.017), and Control vs Stage II+III (p = 6.6 × 10⁻⁶), indicating that co-expression of these immune-fibrotic genes reflects both disease presence and progression. To further examine spatial relationships, we analyzed a recently published spatial transcriptomic dataset that included samples from 6 healthy individuals and 5 IPF patients (https://zenodo.org/records/10012934) [32]. Quantification of spatial interactions using a 20 μm threshold demonstrated that CHI3L1⁺ macrophages and CD44⁺ fibroblasts had significantly higher interaction ratios in IPF lungs compared to controls (Fig. 8D). These findings suggest that close interactions between this macrophage subset and fibroblasts may contribute to the pathogenesis of pulmonary fibrosis.

**Figure 8.**
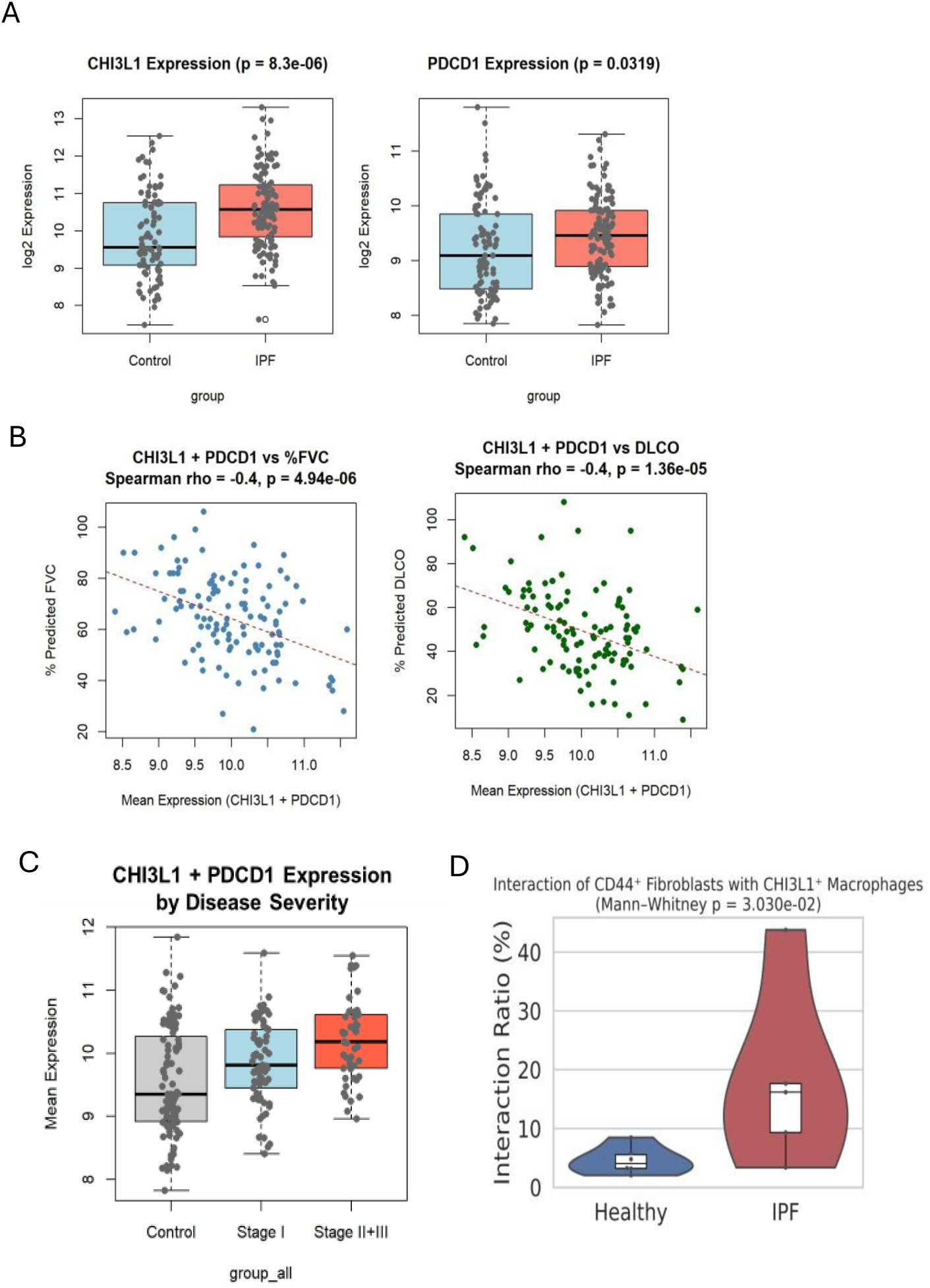
Increased CHI3L1 and PD-1 expression and spatial interaction between CHI3L1⁺ macrophages and CD44⁺ fibroblasts in IPF lungs. (A) Reanalysis of the Lung Tissue Research Consortium microarray dataset (NCBI, GSE47460) demonstrated significantly elevated *CHI3L1* and *PDCD1* (PD-1) expression in IPF lungs compared to controls. (B) Co-expression of *CHI3L1* and *PDCD1* inversely correlated with pulmonary function indices, including % predicted FVC and DLCO. (C) Mean expression of *CHI3L1* and *PDCD1* progressively increased from healthy controls to IPF patients and further stratified by disease severity. Pairwise comparisons showed significant differences between control vs stage I, stage I vs stage II+III, and control vs stage II+III. (D) Analysis of a spatial transcriptomic dataset (6 controls, 5 IPF; https://zenodo.org/records/10012934) revealed significantly higher interaction ratios between CHI3L1⁺ macrophages and CD44⁺ fibroblasts within 20 μm in IPF lungs (*p* < 0.05 by Mann-Whitney U test).

## Discussion

The present study provides compelling evidence that CHI3L1 is a critical mediator of profibrotic macrophage activation and invasive myofibroblast transformation, primarily through CD44-mediated upregulation of PD-L1 expression. Our findings demonstrate that CHI3L1 not only promotes the differentiation of M2 macrophages but also enhances TGF-β- induced invasive properties in myofibroblasts expressing PD-L1. These findings suggest a complex crosstalk facilitated by CHI3L1 that promotes the progression of pulmonary fibrosis, shedding light on potential therapeutic targets for fibrosis intervention.

We further demonstrate that CHI3L1 is both necessary and sufficient to drive alternative (M2) macrophage activation in response to profibrotic cytokines such as IL-13 and TGF-β. Genetic deletion of CHI3L1 markedly reduced the expression of canonical M2 markers (CD206 and CD163) and profibrotic markers (CX3CR1, SiglecF) in bone marrow-derived macrophages and *in vivo* bleomycin models, underscoring its essential role in skewing macrophage phenotype toward a profibrotic state. Moreover, in human THP-1 cells, recombinant CHI3L1 induced M2 differentiation and upregulated PD-L1 expression, a key immune checkpoint molecule known to inhibit T-cell responses and promote fibrosis. These CHI3L1-driven effects were abrogated by either anti-CHI3L1 monoclonal antibody (FRG) or kasugamycin, a pan-chitinase inhibitor, suggesting both therapeutic and mechanistic relevance.

CHI3L1 is expressed at elevated levels in the serum and/or tissues from patients with IPF, and the significant role of CHI3L1 in the pathogenesis of pulmonary fibrosis has been demonstrated in animal models [9]. However, the exact mechanism that CHI3L1 uses to regulate pulmonary fibrosis has not yet been clearly defined. It is interesting to note that CHI3L1 stimulates alternative activation of lung macrophages and mediates the IL-13/TGF-β-induced profibrotic macrophage differentiation, suggesting an essential role of CHI3L1 in pulmonary fibrosis. Consistent with these studies, recent studies using single-cell RNA-seq analysis on the lungs of IPF patients showed CHI3L1 is highly expressed in profibrotic macrophages, called IPF- expanded macrophage (IPFeMϕ), which was predominantly noted in IPF patients [33]. Additional analysis of publicly available single-cell RNA-seq data in other cohorts of patients with IPF or interstitial lung disease (ILD) demonstrated that CHI3L1-expressing macrophages overlapped distinctively with macrophage clusters detected in lungs of IPF patients but not in normal, COPD, or other lung diseases [34–36]. Taken together, these studies suggest that CHI3L1 is a key player driving profibrotic macrophage activation and differentiation in the lungs of IPF patients.

The mechanistic link between CHI3L1 and PD-L1 expression in macrophages represents a key immunoregulatory axis in pulmonary fibrosis. We show that CHI3L1 promotes the differentiation of CD206^+^/CD163^+^ M2 macrophages concomitant with increased PD-L1 expression, and this effect is abrogated by either CHI3L1 neutralization or kasugamycin- mediated inhibition. These findings position PD-L1 as a downstream effector of CHI3L1- driven macrophage polarization. This is consistent with prior studies demonstrating that PD-L1 not only mediates immune suppression but also contributes to the acquisition and maintenance of M2-like phenotypes in macrophages [37, 38], as well as to fibroblast-to-myofibroblast transformation and extracellular matrix remodeling [24, 39, 40]. Thus, the CHI3L1-PD-L1 axis may serve as a central node linking immune regulation with fibrotic progression. Targeting this pathway could simultaneously diminish M2 macrophage-driven immune suppression and fibrogenesis, supporting the therapeutic rationale for CHI3L1 or PD-L1 blockade in fibrotic lung disease.

Beyond macrophage polarization, we found that CHI3L1 potentiates TGF-β-induced myofibroblast transformation, migration, and invasiveness in lung fibroblasts. Mechanistically, CHI3L1 amplified TGF-β-mediated phosphorylation of ERK, AKT, and SMAD2, with ERK activation being particularly dependent on CD44, a known co-receptor of CHI3L1 [28]. Our study further reveals that CD44 is essential for this CHI3L1/TGF-β-induced myofibroblast transformation. CD44 knockdown or neutralization inhibited CHI3L1/TGF-β synergy in promoting α-SMA and PD-L1 expression, as did PD-L1 knockdown. These data identify a novel CHI3L1-CD44-ERK-PD-L1 signaling axis that integrates immunoregulatory and fibrogenic pathways within fibroblasts. This observation aligns with prior research implicating CD44 in pulmonary fibrosis [41–45] and suggests that CD44-PD-L1 axis could be a promising target for inhibiting CHI3L1/TGF-β-stimulated fibroblast activation and fibrosis progression.

In the development and progression of pulmonary fibrosis, the importance of intercellular crosstalk between immune and non-immune cells, including macrophages, epithelial cells and fibroblasts has been well established [30, 46, 47]. Importantly, we observed that CHI3L1 mediates direct crosstalk between profibrotic macrophages and fibroblasts. Macrophages pre-activated with IL-13 and TGF-β induced robust profibrotic effects on fibroblast activation in a CHI3L1- and CD44-dependent manner. This bidirectional loop likely amplifies and sustains the fibrotic cascade, with PD-L1 serving as a shared effector molecule in both cell types. *In vivo*, lungs of CHI3L1-overexpressing transgenic mice exhibited a marked increase in PD-L1^+^ M2 macrophages and PD-L1^+^/PDGFRα^+^ fibroblasts following bleomycin challenge. These changes were accompanied by elevated CD45⁺/PD-1^+^ immune cells, suggesting that CHI3L1 also contributes to immune checkpoint engagement and immune evasion within fibrotic lungs, a phenomenon previously described in cancer biology [21, 27, 48], but less well-characterized in fibrotic disease. Our co-culture experiments provide evidence of CHI3L1-mediated intercellular crosstalk between macrophages and fibroblasts, highlighting a mechanism by which profibrotic macrophages highly expressing CHI3L1 may influence fibroblast activation and contribute to a self-sustaining fibrotic microenvironment. These results support a model in which CHI3L1 facilitates macrophage-fibroblast communication through the CD44-PD-L1 axis, thereby promoting fibroblast activation in response to profibrotic stimuli. This CHI3L1- mediated macrophage-fibroblast crosstalk may represent a critical feedback loop that exacerbates fibrosis, offering potential intervention points to disrupt this cycle leading to persistent and progressive pulmonary fibrosis.

Perhaps most notably, therapeutic blockade of CHI3L1 and PD-1 using bispecific antibodies or combined monotherapies significantly attenuated fibrosis severity, reduced profibrotic macrophage activation, and diminished the expression of fibrotic markers. Treatment with an anti-CHI3L1/PD-1 bispecific antibody was particularly effective, outperforming single-agent therapies. These results provide compelling preclinical evidence that dual targeting of CHI3L1 and PD-1/PD-L1 signaling is a promising strategy for reversing established fibrosis or halting its progression.

In addition to our mechanistic findings *in vitro* and *in vivo*, our analysis of large publicly available human transcriptomic datasets provides important clinical validation. We observed that *CHI3L1* and *PDCD1* (PD-1) transcript levels were significantly elevated in IPF patients compared to controls, and that their co-expression inversely correlated with lung function indices, including FVC and DLCO [31]. Notably, expression progressively increased with disease severity, suggesting that the CHI3L1-PD-1 axis is not only a marker of disease presence but also a driver of disease progression. These observations are consistent with prior reports linking CHI3L1 to IPF severity and extend them by demonstrating its close association with PD-1, a key immune checkpoint molecule that promotes tolerance and fibrotic remodeling [8, 9, 15]. Moreover, spatial transcriptomic analyses demonstrated that CHI3L1⁺ macrophages and CD44⁺ fibroblasts exhibited significantly greater spatial interactions in IPF lungs compared to healthy controls [32]. This finding suggests that CHI3L1 actively facilitates pathogenic macrophage–fibroblast crosstalk within the fibrotic niche, rather than acting solely through diffuse paracrine mechanisms. Together with our co-culture experiments and in vivo studies, these human data support a model in which CHI3L1 orchestrates macrophage-fibroblast communication through a CD44-PD-L1 axis, thereby amplifying TGF-β-driven fibroblast activation and invasive myofibroblast differentiation. Taken together, these human and preclinical data highlight CHI3L1 as a central mediator of immune-fibrotic circuits in IPF. They also strengthen the translational rationale for simultaneously targeting CHI3L1 and PD-1/PD-L1, since such strategies may both relieve macrophage-driven immune suppression and disrupt the pathogenic macrophage-fibroblast interactions that sustain progressive fibrosis

While our study provides compelling evidence supporting the therapeutic value of dual CHI3L1 and PD-1/PD-L1 blockade, several limitations warrant further investigation. Although our *in vitro* and *in vivo* models elucidate the role of CHI3L1 in macrophage-fibroblast crosstalk, the precise molecular mechanisms downstream of PD-L1 that contribute to myofibroblast transformation remain incompletely understood. In particular, dissecting whether PD-L1 functions in its membrane-bound versus soluble form, and how these isoforms interact with PD-1 expressed on macrophages or T cells, will be essential for understanding the full scope of PD-1/PD-L1-mediated immune modulation in fibrotic lung environments. Furthermore, while bispecific antibody therapy targeting CHI3L1 and PD-1 demonstrated enhanced anti-fibrotic efficacy in our preclinical model, comprehensive evaluation of long-term safety, pharmacokinetics, and immunogenicity is required prior to translation into clinical settings. Future studies should include progressive and persistent fibrosis models and expanded immune profiling to better characterize the durability and specificity of the therapeutic response. Ultimately, integrating mechanistic insights with rigorous preclinical testing will be critical for advancing CHI3L1-based immunotherapies for pulmonary fibrosis and related fibrotic diseases.

Taken together, our study defines CHI3L1 as a key pathogenic mediator that bridges immune activation, stromal remodeling, and immune checkpoint signaling in pulmonary fibrosis. These findings expand our understanding of CHI3L1 beyond its known roles in cancer and infection, and offer new therapeutic avenues for fibrotic lung diseases, including idiopathic pulmonary fibrosis (IPF), where current treatment options remain limited. Future studies should explore CHI3L1’s interactions with other immune and stromal cell populations, assess its utility as a biomarker of disease progression, and validate the efficacy of CHI3L1-targeted therapies in diverse models of organ fibrosis.

## Materials and Methods

### Mice and Animal Models

Wild-type (WT), CHI3L1-null mutant (*Chil1*^-/-^), and CHI3L1 Tg mice were generated and characterized in our laboratory as previously described [20, 49]. Experiments were conducted using age (8-10 weeks old) and sex-matched wild-type (WT) and CHI3L1 Tg mice. All protocols for animal experiments were reviewed and approved by the Institutional Animal Care and Use Committee (IACUC) at Brown University

### Bone Marrow-Derived Macrophages (BMDMs)

BMDMs were isolated from the femurs of WT and *Chil1*^-/-^ mice as previously described [50]. Bone marrow was cultured in complete DMEM medium containing 25 ng/mL M-CSF for 7 days to differentiate into macrophages. Macrophages (1 × 10^6^/well in a 6-well plate) were stimulated with recombinant interleukin (rIL)-13 (R&D Systems, 20 ng/mL) and recombinant transforming growth factor (rTGF)-β_1_ (R&D Systems, 5 ng/mL), or recombinant interferon (rIFN)-γ (R&D Systems, 20 ng/mL) for 72 hours and subsequent evaluated.

### Flow cytometric analysis (FACS)

Single-cell suspensions from whole mouse lungs were prepared using the Lung Dissociation Kit (Miltenyi Biotec) according to the manufacturer’s instructions. Cells were stained with fluorescently labeled antibodies against mouse CD163 (Thermo Fisher Scientific, TNKUPJ), mouse CD206 (Thermo Fisher Scientific, MR5D3), mouse CX3CR1 (BioLegend, SA011F11), mouse SiglecF (Thermo Fisher Scientific, 1RNM44N), mouse PD-1 (Thermo Fisher Scientific, J43), mouse PD-L1 (Thermo Fisher Scientific, MIH5), mouse Ly6G (Thermo Fisher Scientific, 1A8-Ly6g), mouse CD45 (BioLegend, 30-F11), human CD163 (BD Bioscience, GHI/61), and human CD206 (BioLegend, 15-2). Flow cytometry data were collected using the BD FACSAria IIIu and the Cytek Aurora and analyzed with FlowJo software (v10). Gating strategy of the macrophages and other cells employed in this study is illustrated in Supplemental Fig. S6.

### THP-1 Cell Differentiation and Macrophage Activation

Human monocytic THP-1 cells (ATCC, TIB-202) were differentiated into macrophages by stimulation with 100 ng/mL phorbol 12-myristate 13-acetate (PMA; Merck) and then rested in complete RPMI-1640 (Thermo Fisher Scientific) for 24 hours. The differentiated cells were then treated with rCHI3L1 (R&D Systems, 500 ng/mL) in the presence or absence of anti-CHI3L1 antibody (FRG; 250 ng/mL) or kasugamycin (KSM; Merck, 250 ng/mL). After 72 hours of incubation, flow cytometry was performed to assess the expression of M2 markers (CD163^+^, CD206^+^), and PD-L1 levels were analyzed by qPCR and Western blotting.

### Double-Label Fluorescent Immunohistochemistry

Formalin-fixed, paraffin-embedded (FFPE) lung tissue blocks were sectioned into 5 µm-thick slices and mounted onto glass slides. Sections were deparaffinized, rehydrated, and subjected to heat-induced epitope retrieval using a steamer and antigen unmasking solution (Abcam, citrate buffer, pH 6.0) for 30 minutes. Blocking was performed with serum-free protein blocking solution (Dako/Agilent, Santa Clara, CA) for 10 minutes at room temperature. Slides were incubated overnight at 4°C with fluorescent-labeled primary antibodies against CD206 (R&D systems, AF2535), CD163 (Abcam, EPR19518), PD-L1 (Invitrogen, #PA5-20343) and PDGFRα (Thermo Fisher Scientific, APA5). After washing, fluorescent-labeled secondary antibodies were applied for 1 hour at room temperature. Sections were counterstained with DAPI and mounted with coverslips.

### RNA Extraction and Semi-Quantitative Real-Time qPCR

Total RNA was isolated using TRIzol reagent (Thermo Fisher Scientific) followed by RNA purification with the RNeasy Mini Kit (Qiagen, Germantown, MD) according to the manufacturer’s instructions. Reverse transcription and real-time PCR were performed as previously described [20]. Ct values for target genes were normalized to internal housekeeping genes *GAPDH* and *Actb*. Primer sequences used in the real-time qPCR are following: *CD274*-S: CAAAGAATTTTGGTTGTGGA, *CD274*-AS: AGCTTCTCCTCTCTCTTGGA; *GAPDH*-S: GAAGGTCGGAGTCAACGGATT, *GAPDH*-AS: CGCTCCTGGAAGATGGTGAT; *Col1a1*-S: GCTCCTCTTAGGGGCCACT, *Col1a1*-AS: CCACGTCTCACCATTGGGG, *Acta2*-S: TCAGCGCCTCCAGTTCCT, *Acta2*-AS: AAAAAAAACCACGAGTAACAAATCAA, *Cd206*-S: CTCTGTTCAGCTATTGGACGC, *Cd206*-AS: TGGCACTCCCAAACATAATTTGA; *CX3CR1*-S: TCTTCACGTTCGGTCTGGTG, *CX3CR1*-AS: AGGATGAGTCTGACGGCTCT; *Actb*-S: GGCTGTATTCCCCTCCATCG, *Actb*-AS: CCAGTTGGTAACAATGCCATGT.

### Western Blotting

Western blots were used according to standard protocols as previously described [21]. Antibodies against pSMAD2 (Cell Signaling Technology (CST), 138D4), p-ERK (CST, Polyclonal), p-AKT (CST, 193H12) and α-SMA(Abcam, EPR5368), Smad2 (CST, L16D3), Erk (CST, Polyclonal) Akt (CST, 11E7), CD44, (CST, E7K2Y), CD163 (Invitrogen, Polyclonal), CD206 (Invitrogen, JF0953), PD-L1 (CST, E1L3N), β-Actin (Santa Cruz, C4), β-Tubulin (CST, #2146) and mouse PD-L1 (CST, D4H1Z).

### Fibroblast Migration Assays

NHLFs were pre-treated with rTGF-β (5 ng/mL), rCHI3L1 (500 ng/mL), FRG, or KSM for 48 hours. Following stimulation, 3 × 10^4^ cells were seeded into the upper chamber of 24-well transwell insert with 8 μm pore membranes (Corning, #353097) in serum-free DMEM. The lower chamber was filled with DMEM containing 10% FBS as a chemoattractant. After 24 hours, non-migrating cells on the upper surface of membrane were removed using a cotton swab. Migrated cells on the lower surface were stained with Kwik-Diff™ solution (Thermo Fisher Scientific) and subsequently quantified.

### Fibroblast Invasion Assays

NHLF pre-treated described in the migration assay. Subsequently, 8 × 10^4^ cells were seeded into 8 μm pore BioCoat™ Matrigel invasion chambers (Corning, #354480) in serum-free DMEM. The lower chamber was filled with serum-free DMEM containing recombinant platelet-derived growth factor-BB (rPDGF-BB, 10 ng/mL; PeproTech, #100-14B) as a chemoattractant. After 24 hours, non-invading cells on the upper surface of membrane were removed. Invading cells on the lower surface were stained with Kwik-Diff™ solution and subsequently quantified.

### Primary Lung fibroblasts and siRNA Silencing

Murine lung fibroblasts were isolated and cultured according to the reported procedures [51]. NHLF or primary lung fibroblasts were transfected with siRNA using Lipofectamine RNAiMAX (Thermo Fisher Scientific). siRNA targeting human PD-L1 (100 nM; Bioneer, #29126-1 and #29126-2) and mouse CD44 (100 nM; Bioneer, #12505-1 and #12505-2) or control siRNA (100 nM; Bioneer, #SN-1003) were used. The efficiency of PD-L1 or CD44 silencing was evaluated by Western blot analysis.

### Co-Culture Studies

Co-culture experiments were performed to assess the interaction between macrophages and fibroblasts. WT or *Chil1^-^*^/*-*^ BMDMs were stimulated with rIL-13 (20 ng/mL) and rTGF-β (5 ng/mL) for 72 hours. Primary lung fibroblasts were then co-cultured with macrophages, and after 48 hours, the expression of α-SMA and PD-L1 in the fibroblasts was analyzed by Western blot. To evaluate the effect of CD44 on fibroblasts from stimulated macrophages, CD44- silenced and control fibroblasts were co-incubated with TGF-β/IL-13-stimulated WT macrophages for 48 hours.

### Bleomycin-Induced Pulmonary Fibrosis and Histological Analysis

To assess the role of CHI3L1 in lung fibrosis, WT and CHI3L1 Tg mice were given doxycycline (Merck, #D9891) in drinking water for 7 days, followed by intratracheal injection of bleomycin (2.5 U/kg body weight). 14 days after bleomycin administration, lung tissues were collected for FACS or histological analysis. Hematoxylin and eosin (H&E) staining was performed to assess lung architecture, and Masson’s trichrome staining was used to assess collagen deposition. Sircol collagen assay was used to measure total collagen accumulation in the lung as previously described [20].

### Therapeutic Treatment with Antibodies

To evaluate the therapeutic potential of targeting CHI3L1 and the PD-1/PD-L1 axis in pulmonary fibrosis, mice were intraperitoneally injected with either monotherapies (anti-PD-1 or anti-CHI3L1 antibody [FRG], each at 8 mg/kg), a combination of both antibodies (8 mg/kg each), or a bispecific anti-CHI3L1×PD-1 antibody (8 mg/kg). Antibody treatments were administered every other day from day 7 to day 14 following bleomycin challenge. Lung fibrosis was evaluated by histological analysis, Masson’s trichrome staining, and gene expression of fibrosis markers (*Col1a1* and *Acta2*) and profibrotic macrophage markers (*Cd206* and *Cx3cr1*). FACS analysis was used to assess the activation of profibrotic macrophages.

### Spatial Interaction Analysis

Spatial transcriptomics data from [32] were analyzed to evaluate potential physical interactions between CHI3L1⁺ macrophages and CD44⁺ fibroblasts. Macrophages were defined based on cell-type annotation and classified as CHI3L1⁺ if CHI3L1 expression exceeded a predefined threshold (1.0), empirically determined from the distribution of expression values. Similarly, fibroblasts were classified as CD44⁺ if annotated as fibroblasts and expressing CD44 above 1.5 normalized units. For each sample, the spatial coordinates of CHI3L1⁺ macrophages and CD44⁺ fibroblasts were extracted. Using a k-d tree algorithm, we computed the nearest-neighbor distances from each CHI3L1⁺ macrophage to the closest CD44⁺ fibroblast. Cells were considered “interacting” if the distance was ≤ 20 μm, consistent with previous spatial transcriptomics studies [52, 53]. The interaction ratio was defined as the proportion of CD44⁺ fibroblasts that were spatially proximal (≤ 20 μm) to any CHI3L1⁺ macrophage. Interaction ratios were compared between IPF and healthy samples using the Mann-Whitney U test. Additionally, all pairwise distances were aggregated and subjected to the Mann-Whitney U test for group-level comparisons. Such nearest-neighbor and distance-threshold based proximity analyses are widely used in spatial transcriptomics studies [54, 55]. All spatial analyses were performed in Python.

### Statistical Evaluations

All statistical analyses were conducted using GraphPad Prism software. Comparisons between groups were made using two-tailed Student’s *t* tests. For multiple group comparisons, one-way ANOVA followed by appropriate post hoc tests was performed. Mann-Whitney U test was used to test the distributions of two independent samples when the data were not normally distributed or were at least ordinal. Data are presented as mean ± SEM, and statistical significance was defined as *p* < 0.05.

## Acknowledgements

This work was supported by the National Institutes of Health (NIH) grants PO1 HL114501 (J.A.E.), R01HL155558 (C.G.L.), T32 HL134625 (T.S.) and Department of Defense grant W81XWH-22-1-0041 (C.G.L.). This work was partially supported by the National Research Foundation (NRF) grant funded by the Korea government (MSIT) (#RS-2024-00405542) (S.J.S. and C.G.L.).

## Author contributions

C.G.L. conceptualized and designed the study. H.S.J., T.S., J.H.L., S.K., B.M., and Y.Z. performed experiments and analyzed data. H.S.J., T.S. contributed to data curation and statistical analysis. C.G.L. H.S.J. and T.S. wrote the original draft. C.G.L. supervised the overall experimental process. J.A.E. and S.J.S. provided critical comments on the experimental data. All authors reviewed and edited the manuscript and approved the final version for submission

## FIGURES AND FIGURE LEGENDS

**Figure S1.**
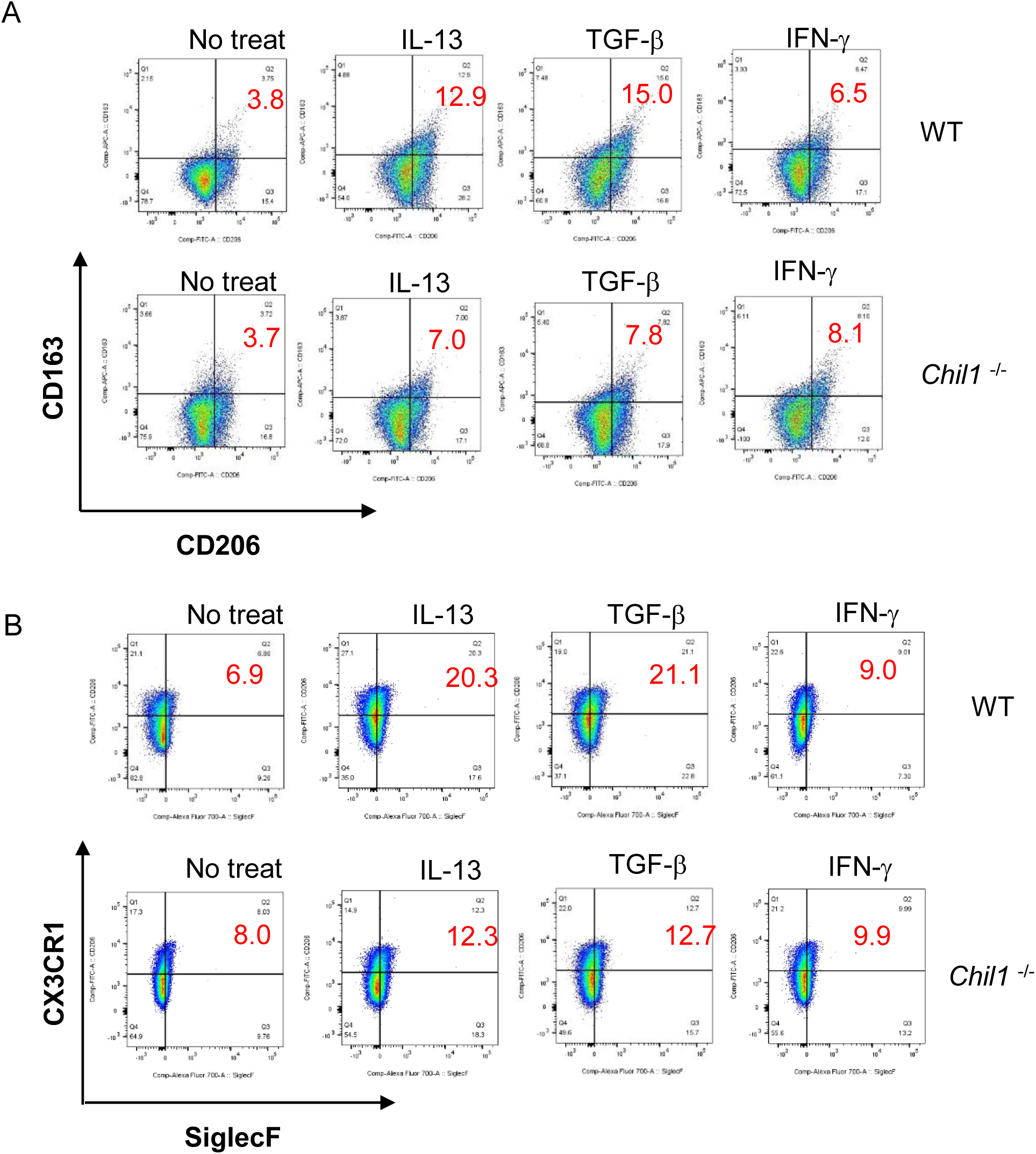
CHI3L1 promotes M2 macrophage polarization in response to IL-13 or TGF-β. Bone marrow-derived macrophages (BMDMs) were isolated from wild-type (WT) and CHI3L1-null (*Chil1*^-/-^) mice and stimulated with recombinant IL-13 (20 ng/mL), TGF-β_1_ (5 ng/mL), and IFN-γ (20 ng/mL) separately for 72 hours. Cells were then analyzed by flow cytometry. (A) Frequency of CD206^+^/CD163^+^ M2 macrophages. (B) Frequency of CX3CR1^+^/SiglecF^+^ profibrotic macrophages.

**Figure S2.**
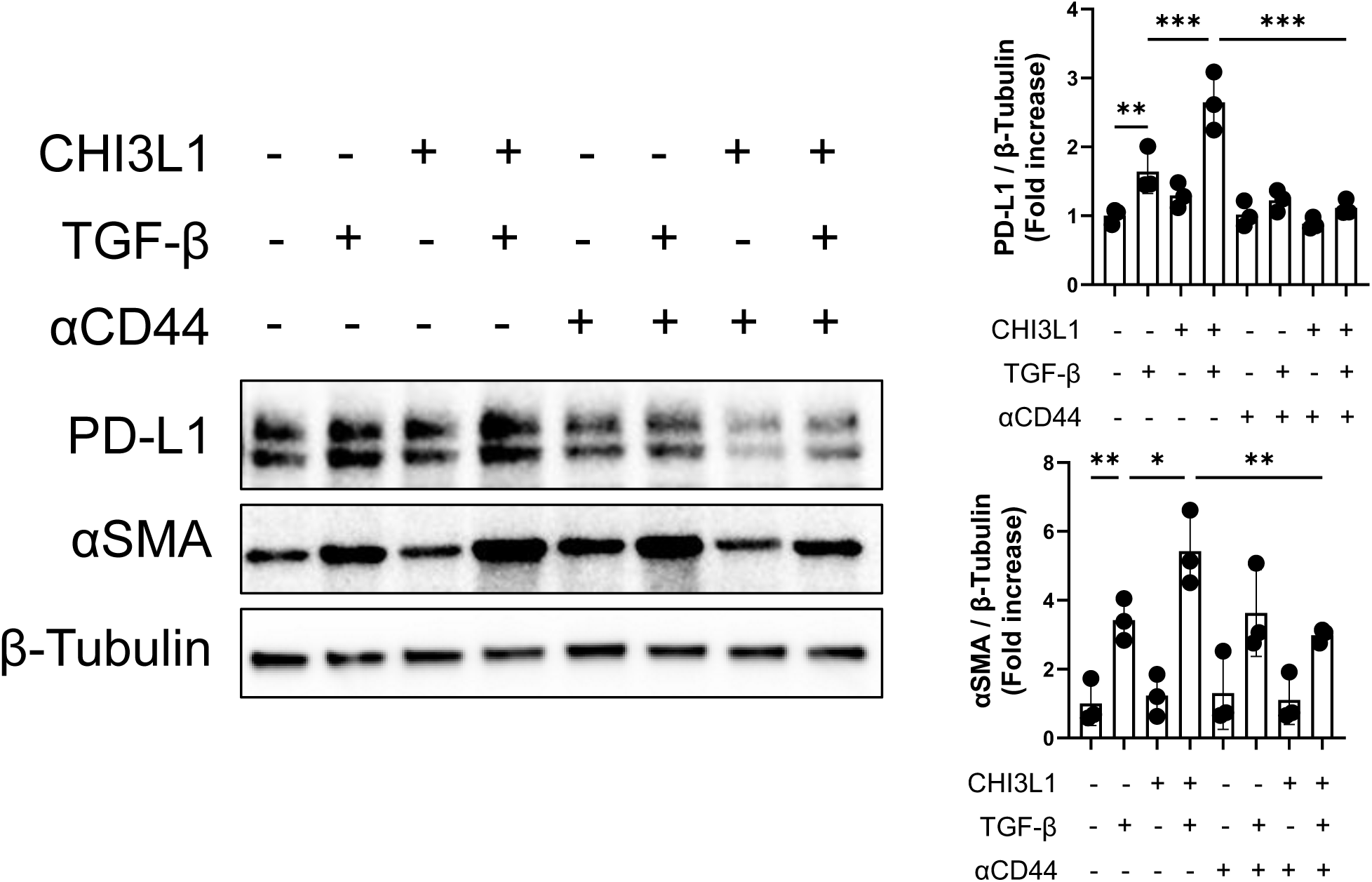
Role of CD44 in CHI3L1/TGF-β stimulated expression of α-SMA and PD-L1 expression in lung fibroblasts. Western blot analysis of PD-L1 and α-SMA expression in lung fibroblasts stimulated with recombinant CHI3L1 (500 ng/mL) and/or TGF-β_1_ (5 ng/mL) in the presence or absence of neutralizing anti-CD44 antibody (Thermo Fisher Scientific, 5 μg/mL). Bar graphs represent densitometric quantification of protein expression (mean ± SEM). \**p* < 0.05, \*\**p* < 0.01, \*\*\**p* < 0.001 by one-way ANOVA with multiple comparisons.

**Figure S3.**
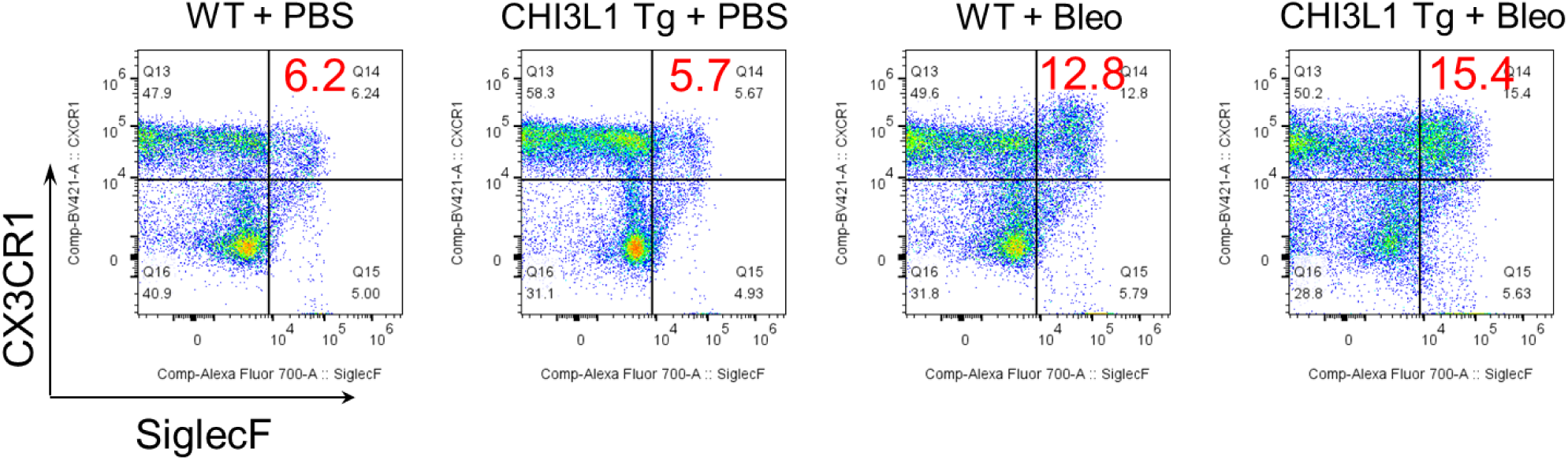
CHI3L1 increases profibrotic macrophage activation in bleomycin-challenged lungs. Six to eight weeks old WT and CHI3L1 Tg mice were challenged with PBS or bleomycin then the mice were sacrificed on 14 days after bleomycin challenge and subjected to the analysis. (A) Flow cytometric evaluations on CX3CR1^+^ and Siglec F^+^ macrophages in lungs of WT and CHI3L1 (YKL-40) transgenic (Tg) mice.

**Figure S4.**
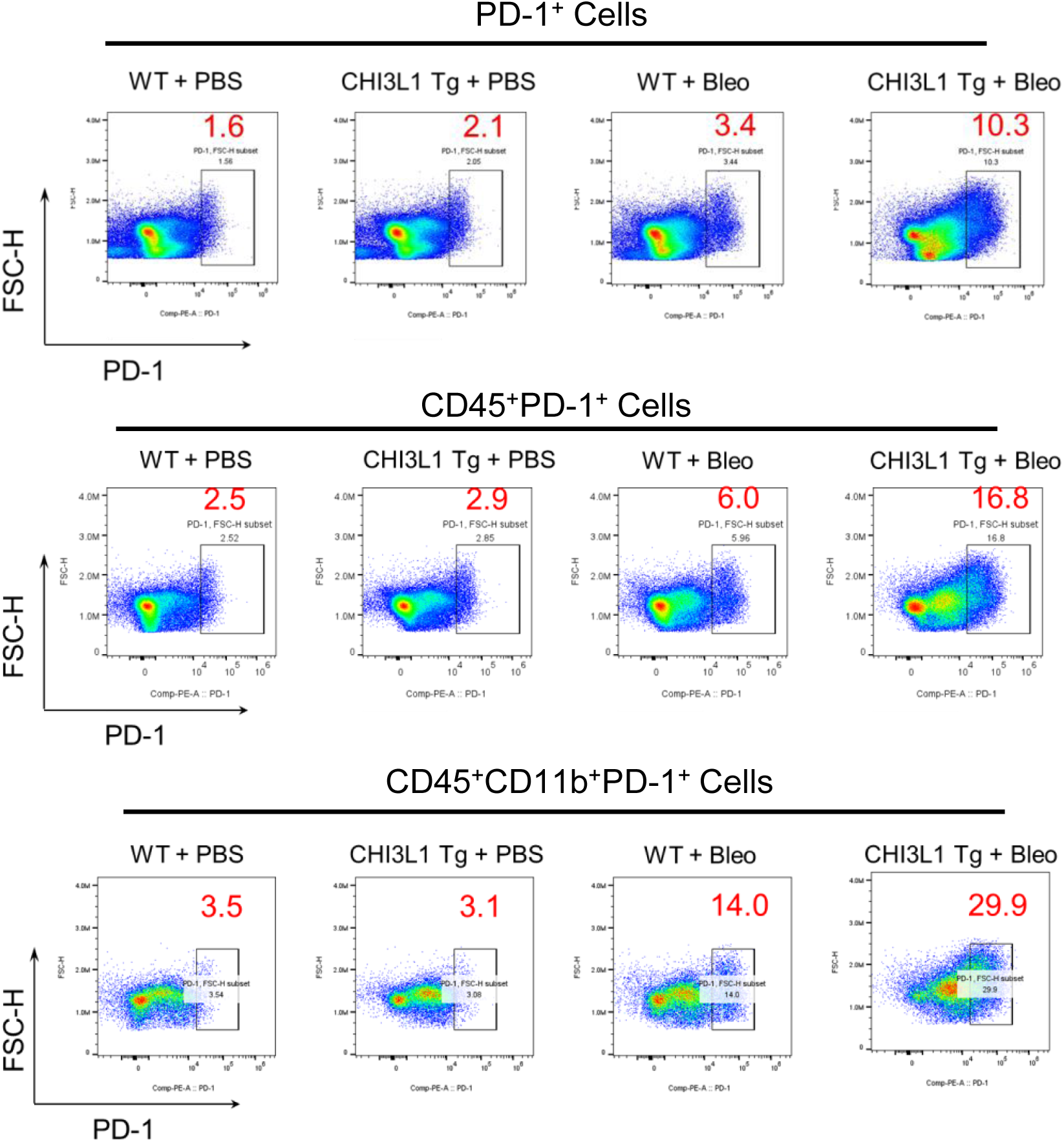
CHI3L1 increases total and PD-1–expressing inflammatory cells in bleomycin-challenged lungs. Six to eight weeks old WT and CHI3L1 Tg mice were challenged with PBS or bleomycin then the mice were sacrificed on 14 days later for analysis. Flow cytometric analysis demonstrated increased numbers of total inflammatory cells and CD45⁺/PD-1⁺ cells, including CD45⁺/CD11b⁺/PD-1⁺ macrophages, in the lungs of CHI3L1 Tg mice compared with WT controls.

**Fig. S5.**
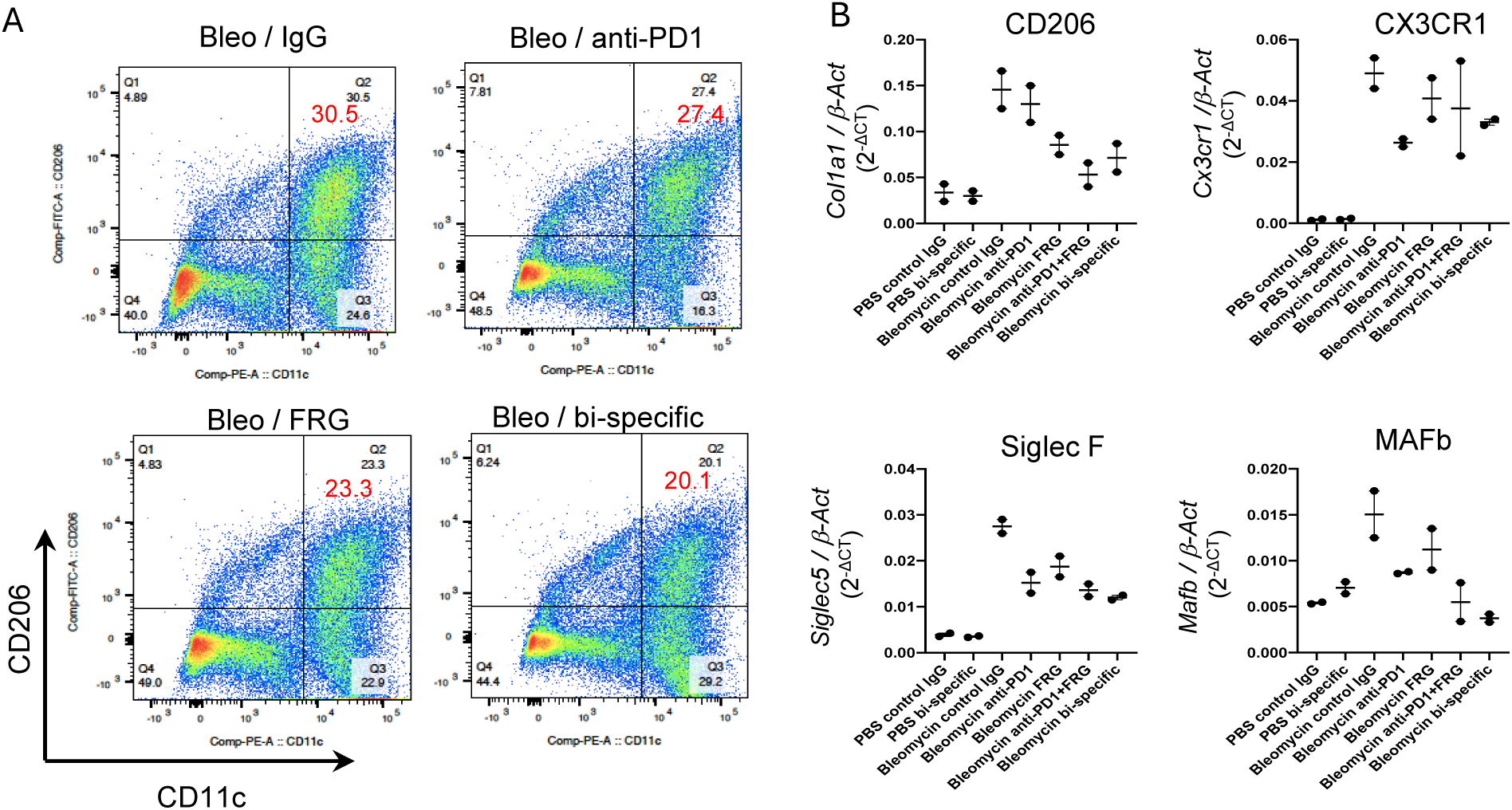
Regulation of profibrotic macrophage activation in bleomycin challenged lungs by mono-and bi-specific anti-CHI3L1 antibodies. (A) FACS analysis of CD45^+^/F4/80^+^/CD11c^+^/CD206^+^ profibrotic macrophages. (B) qRT-PCR analysis of the mRNA expression of *Cd206*, *Cx3cr1*, *Siglec5* and *Mafb* in sorted CD45^+^/F4/80^+^/CD11c^+^ macrophages following antibody treatments.

**Figure S6.**
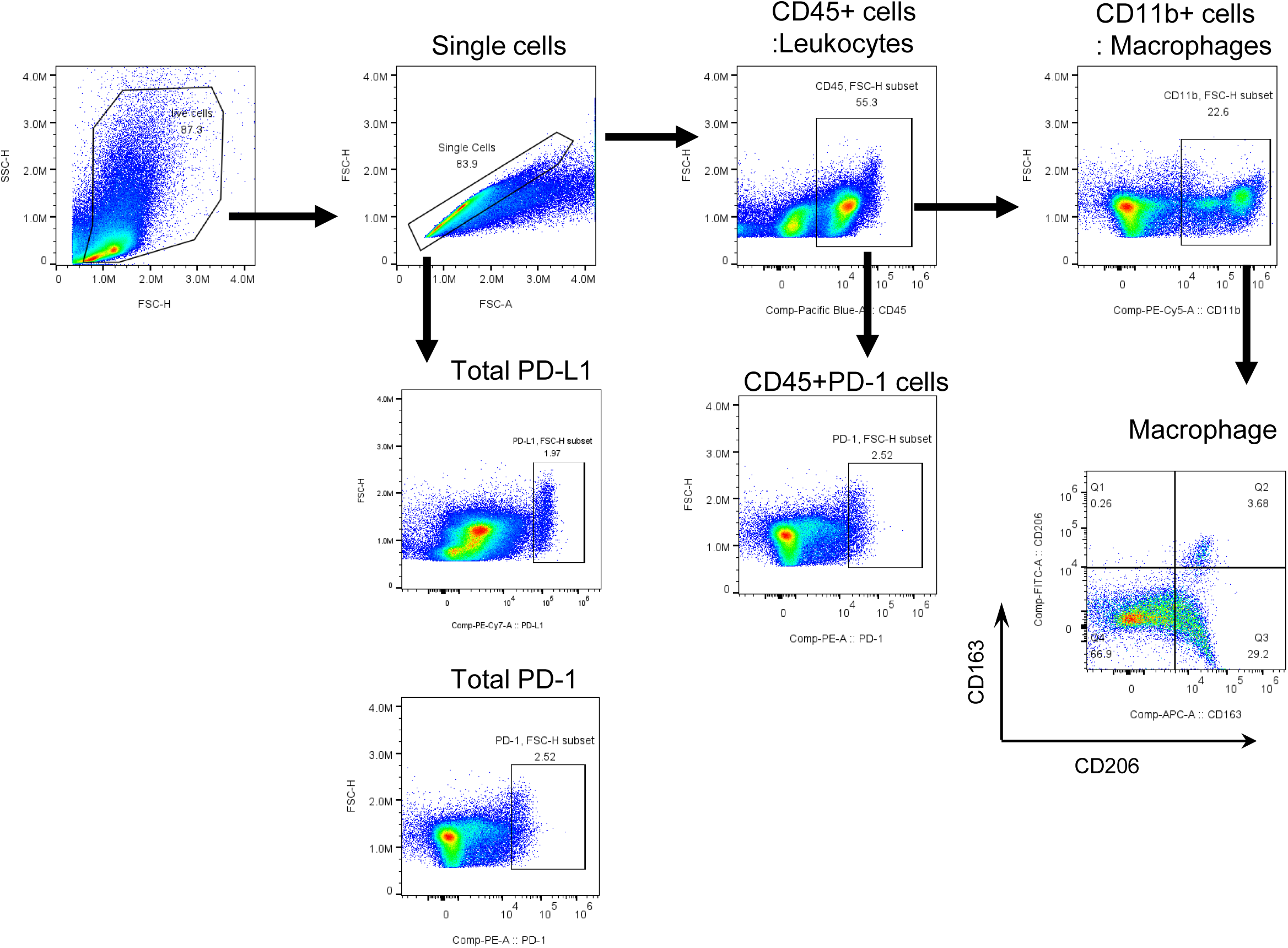
Gating strategies used for flow cytometric in this study.

